# Stomatal dynamics are regulated by leaf hydraulic traits and guard cell anatomy in nine true mangrove species

**DOI:** 10.1101/2023.02.01.526604

**Authors:** Ya-Dong Qie, Qi-Wei Zhang, Scott A. M. McAdam, Kun-Fang Cao

## Abstract

Stomatal regulation is critical for mangroves to survive water deficits and highly fluctuating ambient water availability in the hyper-saline intertidal zone. Despite the importance of stomatal regulation in mangroves very little is known about stomatal sensitivity to vapour pressure deficit (VPD), and the co-ordination of this trait with stomatal morphology and leaf hydraulic traits in these species.

We measured the stomatal response to a step increase in vapour pressure deficit (VPD) *in situ*, stomatal anatomy, leaf hydraulic vulnerability and pressure-volume traits in nine true mangrove species of five families. We aimed to answer two questions: (1) Does stomatal morphology determine stomatal dynamics in response to a high VPD in mangroves and (2) do leaf hydraulic traits influence stomatal sensitivity to VPD in mangroves?

We found that the stomata of mangrove plants highly sensitive to VPD, and that species with higher maximum stomatal conductance had slower stomatal responses to an increase in VPD, and that stomatal density and size were correlated with the speed of stomatal closure at high VPD across the closely-related species. We also found that a higher leaf capacitance (*C*_leaf_) and more resistance to leaf hydraulic vulnerability were associated with slower stomatal responses to an increase in VPD.

Our results demonstrate that the dynamics of the stomatal response to an increase in VPD are regulated by leaf hydraulic traits and stomatal morphology. Our work provides a quantitative framework to better understand stomatal regulation in mangroves in an environment with highly dynamic water availability.

## Introduction

Stomata on the leaves of vascular plants dynamically control transpirational water loss (Meinzer, 1993). Under sufficient soil moisture and saturating light, stomatal conductance to water vapour (*g*_s_) is mainly regulated by vapor pressure difference between leaves and the atmosphere (VPD). Stomata open to maximal extent at a low VPD in the morning, achieving the highest rates of carbon assimilation at this time. Once VPD begins to increase as the day progresses stomata tend to close, reducing water potential declines to avoid a critical water tension that would induce xylem embolism (Sperry, 2000; Brodribb & Holbrook, 2004a; Choat *et al*., 2018; Durand *et al*., 2019). In angiosperms, after a sudden increase in VPD, stomata often open instantaneously because of a passive contraction of the epidermal cells, termed the “wrong-way” response (WWR); following a period of opening stomata then close gradually and reach a new final steady-state (Buckley, 2016). This stomatal closure after wrong-way opening is believed to be driven by the hormone abscisic acid (ABA), which is synthesized when mesophyll cells approach turgor loss point and closes stomata (McAdam & Brodribb, 2016). Unlike in species of lycophyte, fern and conifer, in which the kinetics of stomatal responses to VPD can be readily predicted by a passive hydraulic model of stomatal regulation (McAdam & Brodribb, 2015), in angiosperms stomatal closure at high VPD is believed to be largely driven by an active metabolic signal like ABA, and the kinetics may not be easily predictable from anatomical traits. Stomatal kinetics from initial state to final homeostasis in response to environmental fluctuation influences the balance between carbon assimilation (*A*) and water loss and for this reason differences in kinetics can be adaptively relevant (Buckley, 2016; Lawson & Vialet-Chabrand, 2019; Durand *et al*., 2019; Grossiord *et al*., 2020). The fast stomatal kinetics, by closely tracking environmental disturbances, could enable stomata to operate optimally (Cowan & Farquhar, 1977; Drake *et al*., 2013; Anderegg *et al*., 2018; Eyland *et al*., 2021). Moreover, fast stomatal closure restricts transpirational water consumption and prevents hydraulic failure events during rapid changes in high atmospheric water deficit (Brodribb & Holbrook, 2004b; Martin-StPaul *et al*., 2017; Rodriguez-Dominguez & Brodribb, 2020; Carminati & Javaux, 2020).

The rapidity of the *g*_s_ response to an increase in VPD in angiosperms might arise mechanistically from stomatal morphology (e.g., stomatal size and density). Smaller stomata have faster kinetics and greater responses to change in light, owing to a greater guard cell membrane surface area to volume ratio, which facilitates more rapid ion exchange (Drake *et al*., 2013). This hypothesis has been supported by some studies (Drake *et al*., 2013; Raven, 2014; Xiong *et al*., 2018; Kardiman & Ræbild, 2018; Durand *et al*., 2019), but has not yet reached consensus (Elliott-Kingston *et al*., 2016; McAusland *et al*., 2016; Lawson & Vialet-Chabrand, 2019). Furthermore, species with smaller, denser stomata may require less ion uptake or release to establish sufficient osmotic pressure to inflate or deflate against the pressure of the surrounding epidermal pavement cells (DeMichele & Sharpe, 1973; Franks, 2003; D’Ario & Sablowski, 2019). This is associated with a higher bulk leaf osmotic potential at full turgor (π_o_) (Nonami & Schulze, 1989) and thus achieving a higher *g*_max_ (Franks, 2006; Henry *et al*., 2019). However, some studies have not found a relationship between π_o_ and *g*_max_ (Farrell *et al*., 2017; Li *et al*., 2018).

A high *g*_max_ in angiosperms inevitably accelerates transpiratory water loss (Brodribb *et al*., 2007; McAusland *et al*., 2016), which is supplied by an increased investment in vein density to achieve high water transport efficiency (Brodribb *et al*., 2005; Sack & Frole, 2006; Brodribb *et al*., 2007), while maintaining the balance between water supply and evaporative demand (Buckley, 2005; Sack & Holbrook, 2006; Brodribb & Jordan, 2008). Recent studies have demonstrated tight coordination across terrestrial plants between maximum leaf hydraulic conductance (*K*_leaf_) and *g*_max_ (Brodribb *et al*., 2005; Sack & Holbrook, 2006; Zhang *et al*., 2015; Xiong & Nadal, 2020). Leaf hydraulic conductance and *g*_s_ can be decoupled from stomatal responses to temporal rise in VPD due to hydraulic feedbacks that depend on leaf hydraulic resistance and water status (Buckley, 2005; Buckley, 2019). During drought-induced dehydration, leaf hydraulic conductance often declines rapidly, which can further drive stomatal closure as a consequence of decrease of leaf water potential (Ψ_leaf_) (Brodribb *et al*., 2014; Brodribb *et al*., 2016; Skelton *et al*., 2018; Scoffoni *et al*., 2018). By contrast, in some species the leaf hydraulic system has strong resistance to water stress, delaying stomatal closure (Scoffoni *et al*., 2012; Skelton *et al*., 2017; Scoffoni *et al*., 2017; Scoffoni *et al*., 2018). The extent of the water potential of inducing loss in leaf hydraulic conductance and then regulating stomatal closure is still unknown. Stomatal regulation is controlled also by leaf water content (*WC*_leaf_) with changing in cell volume, and hence the cell turgor pressure (Trueba *et al*., 2019; Fu *et al*., 2019; Xiong & Nadal, 2020). Leaf capacitance (*C*_leaf_) is considered a key trait for defining pools of water in leaves alleviating against rapid fluctuations in water potential and being involved in desiccation avoidance (Schulte, 1992; Scholz *et al*., 2011; Xiong & Nadal, 2020). Stomatal closure may thus be delayed in plants with a high *C*_leaf_. Outside ferns and lycophytes (Martins *et al*., 2016) the relationship between the speed of stomatal closure and *C*_leaf_ has been rarely examined explicitly.

Mangrove plants live in intertidal coastal zones of the tropics and subtropics (Duke *et al*., 1998), an environment that is associated with high salt concentration and hypoxia in the soil rhizosphere as well as high atmospheric evaporative demand (Reef & Lovelock, 2015). To adapt to such habitats, mangroves have developed efficient regulation of water consumption, high resistance to xylem embolism, high tolerance to desiccation and high leaf capacitance (Reef & Lovelock, 2015; Jiang *et al*., 2017; Jiang *et al*., 2021; Jiang *et al*., 2022; Aritsara *et al*., 2022). Mangroves have longer guard cell length for a given stomatal density than other vascular plants (Agduma *et al*., 2022). Mangroves are confronted with increasing challenges for survival from global warming, which will result in dramatically increase in VPD, changes that may have led to dramatic die-off events in mangrove forests (Gauthey *et al*., 2022). Yet, relatively little is known about the stomatal dynamics of mangroves in response to high VPD, information that would help us to understand the physiological response of mangroves following any hydraulic perturbation. In this study, we quantified the speed of stomatal closure in response to a step-rise in VPD *in situ*, and performed measurements of stomatal morphological characteristics, pressure–volume curves and hydraulic vulnerability to dehydration of leaves of nine true mangrove tree species. We aimed to answer the following two questions: (1) Does stomatal morphology determine stomatal dynamics in response to a high VPD in mangroves? (2) Do leaf hydraulic traits influence stomatal sensitivity to VPD in mangroves?

## Materials and Methods

### Study site and plant material

The study sites were located at the Dongzhai Harbor National Nature Reserve in Hainan (19°57 ‘N, 110°35’ E), which has the largest mangrove forest area in China, and the Qinglan Harbor Provincial Nature Reserve in Hainan (19°37’ N, 110°50’ E), which has the highest richness of mangrove species in China, respectively. The two mangrove reserves are about 60 km apart and have similar climates, classified as a maritime monsoonal climate. Mean annual temperatures are 23.8 °C and 23.9 °C for the two sites, respectively; mean annual precipitation is about 1700 mm, which is largely concentrated from May to November; mean annual evaporation is about 1800 mm; mean annual relative humidity is 85% and 87%, respectively. The soil types are mainly sandy saline soil and marsh saline soil; the salinity of soil pore water is 3-35 ‰ (Zhang *et al*., 2011) and 10-34 ‰ (Zheng, 2019; Leng, 2020), respectively.

Five mangrove species were selected in Dongzhai Harbor, including *Aegiceras corniculatum, Avicennia marina, Ceriops tagal, Kandelia obovata*, and *Rhizophora stylosa*. Four mangrove species were selected in Qinglan Harbor, including *Bruguiera gymnorhiza, B. sexangula, Sonneratia alba*, and *Xylocarpus granatum*. These species are the dominant or common species in the communities. They vary considerably in leaf structure and stomatal morphology (Chen, 2020), and are thus we would expect if anatomy was regulating stomatal responses in these species to find a diversity of stomatal regulation strategies. Three to six mature, healthy and sun-exposed individuals of each species were selected for the experimental measurements. The information about height, basal diameter (BD) or breast-height diameter (DBH), and the number of cell layers of the leaf hypodermis of the sampled plants are shown in Table 1.

**Table 1.** The basic information for the sample plants of nine mangrove species studied in Hainan, China. *Avicennia marina* only has hypodermal layers beneath the upper epidermis (marked with an asterisk), the rest species have hypodermal layers beneath both upper and lower epidermises. DBH, diameter at breast height, BD, basal diameter.

| Species | Abbr. | Family | Mean Height<br>(range) (m) | Mean DBH/BD<br>(range) (cm) | Number of<br>Hypodermis |
| --- | --- | --- | --- | --- | --- |
| *Aegiceras corniculatum* | AC | Myrsinaceae | 2.1(1.8, 2.4) | 4.8(4, 6) | 4-6 |
| *Avicennia marina* | AM | Verbenaceae | 2.8(2.6, 3) | 7.2(6.7, 8.2) | 6-8* |
| *Bruguiera gymnorhiza* | BG | Rhizophoraceae | 4.7(4, 6.8) | 18.1(11.5, 22.4) | 2 |
| *Bruguiera sexangula* | BS | Rhizophoraceae | 5.8(4.4, 7) | 18.9(10, 29) | 2 |
| *Ceriops tagal* | CT | Rhizophoraceae | 2.3(2.1, 2.5) | 9.2(7.5, 11.8) | 3-4 |
| *Kandelia obovata* | KO | Rhizophoraceae | 3.7(3.2, 4.1) | 10.3(8, 14) | 4 |
| *Rhizophora stylosa* | RS | Rhizophoraceae | 4(3.6, 4.4) | 7.6(6.7, 9.5) | 6-9 |
| *Sonneratia alba* | SA | Sonneratiaceae | 8(6.5, 8.9) | 22.66(14, 33.4) | 0 |
| *Xylocarpus granatum* | XG | Meliaceae | 4.7(4, 5.5) | 18.76(11, 27.5) | 3-5 |

### Kinetics of stomatal response to temporal rise in VPD

Between August and October 2020, one fully expanded leaf from the sun-exposed canopy of each individual tree in the field was selected for gas exchange measurement using a portable photosynthesis system (Li–Cor 6800; Li–Cor Inc.) on a clear day between 08:00 and 11:00 h (local time). Leaves were acclimatized inside the leaf cuvette (Photosynthetic Active Radiation, PAR, 1600 μmol·m^-2^·s^-1^; CO_2_ concentration, 400 ppm; relative humidity, RH, 70%; flow rate, 500 μmol·s^-1^) until stomatal conductance (*g*_s_) reached an initial steady-state (*SS*_initial_; defined as a variation < 5% over 3 min), during which the leaf chamber temperature were controlled to 0-2 °C below the air temperature. At this point, RH was then switched to 20%, resulting in a step-increase in VPD while no change in other environmental parameters, and *g*_s_ showed a transient increase, called the “wrong-way” response (WWR), then declined gradually until it reached a new final steady-state (*SS*_final_). The VPD was about 0.95 kPa and 2.65 kPa before and after changing RH, respectively. The value of *g*_s_ was recorded every 20 s during the entire measurement period. After the measurements of gas exchange, the intact branches from which the leaves grew were marked for subsequent measurements of pressure–volume curves and hydraulic vulnerability to dehydration of leaves.

### Stomatal morphology

Leaves used for gas exchange measurements were collected, enclosed in plastic bags, and immediately stored into the fridge at 4 °C. Paradermal section were sliced from each leaf with a square puncher (∼1 cm^2^) and soaked overnight in a bleach solution composed of hydrogen peroxide and acetic acid (H_2_O_2_:CH_2_COOH=1:1) at 70 °C in an oven. Transparent epidermal samples were isolated from the mesophyll and washed with distilled water. These samples were stained with Safranin-Alcian blue and then mounted on microscope slides. They were observed under a light microscope (Leica DM 3000 LED Wetzlar Germany), capturing five images per leaf at magnifications of 10× for stomatal density and 40× for stomatal size measurements. ImageJ software was used to measure stomatal number and guard cell length and width. Stomatal density (*SD*) was the number of stomata per mm^2^, and theoretical maximum stomatal conductance (*G*_s-max_) was calculated according to the method described by de Boer *et al*. (2016).

### Modelling *g*_s_ responses to a step-rise in VPD

Stomatal temporal dynamics in response to a single step-rise in VPD was fitted empirically using an analytical sigmoidal model (Vialet-Chabrand *et al*., 2013) to evaluate the specific parameters in the response curve of *g*_s_ (Fig. 1). It describes as (Eqn 1):

**Figure 1.**
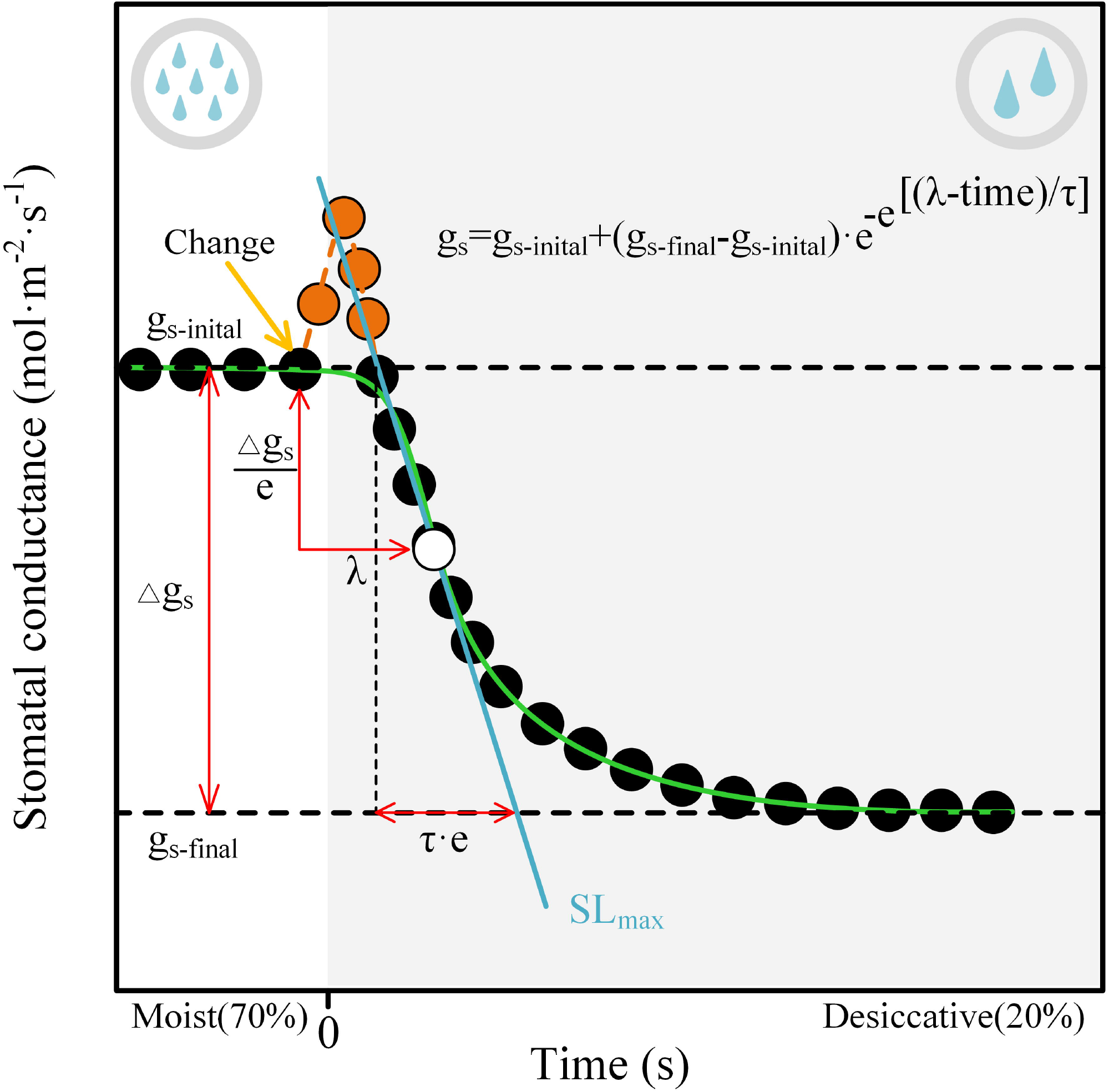
Summary of the parameters derived from an analytical sigmoidal model of changes in stomatal conductance (*g*_s_) in response to a single step-rising vapour pressure deficit (VPD). The yellow arrow showed the time at which vapour-pressure deficit was increased. *g*_s-initial_ and *g*_s-final_ are the stomatal conductance values at the initial and the final steps of the curve, respectively. Δ*g*_s_ is the magnitude of changes in *g*_s_ (Δ*g*_s_ = *g*_s-initial_ - *g*_s-final_). λ is the time between the VPD change and the moment where the change of *g*_s_ is at a maximum (white dot). *SL*_max_ is the tangent (blue line) that goes through this point and is determined as Δ*g*_s_/(τ·e) where e ≈ 2.718. The orange points represent stomatal “wrong-way” response (WWR) that a transient passive opening due to a rapid reduce in epidermal turgor and a less backpressure on the guard cells results from rising transpiration following elevating VPD. Our goal was to discuss the effect of hydraulic-related traits on stomatal VPD response time and thus curve-fitting (green line) without inclusion of data of WWR. The blank area indicates a moist status with relative air humidity of 70%, and while grey area indicates a desiccative status with relative air humidity of 20%.

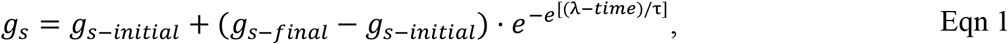

where *g*_s_ (mol·m^-2^·s^-1^) is the *g*_s_ at the corresponding time (s), λ the initial time lag of the sigmoidal curve (s), τ the time constant of *g*_s_ response (s), *g*_s-initial_ and *g*_s-final_ (mol·m^-2^·s^-1^) are the steady-state values of *g*_s_ at the initial and final stages of a sigmoidal curve, respectively. *e* is Euler’s number (c. 2.718). Based on these parameters, we obtained the second parameter, the maximum slope (*SL*_max_) (mmol·m^-2^·s^-1^) (Eqn 2) as an estimator of combining speed and amplitude of the *g*_s_ response to the step-rise in VPD:

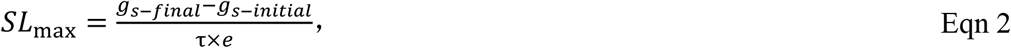

More details on the model and its parameters can be referred to the literature (Vialet-Chabrand *et al*., 2013; Gérardin *et a*l., 2018; Durand *et al*., 2019).

### Pressure–volume curves

Pressure–volume measurements were performed on leaves using the bench drying procedures described by Tyree & Hammel (1972). A branch labeled on the gas exchange measurement was harvested for each individual and rehydrated overnight for next-day measuring. One fully expanded leaf was collected from each rehydrated branch and dehydrated on the bench in a temperature-controlled room with a good-ventilation. The leaf was repetitively weighted and measured for water potential with a pressure chamber (PMS Instruments). The early and late interval time of each curve measurement was 0.5-2h and 4-8h, respectively. Determining a complete pressure– volume curve with least 10 points required leaves to dehydrate for 2 to 4 days. Furthermore, Leaf area was analysed using a scanner and dry mass was determined after oven-drying for 72 h at 60 °C. From the pressure –volume curves, the following parameters were determined: osmotic potential at full turgor (π_o_) and at turgor loss point (π_tlp_), bulk modulus of elasticity (ε) and leaf area specific capacitance at full turgor (*C*_leaf_).

### Leaf hydraulic vulnerability curves

Branches, approximately 110 cm in length, neighboring those measured for gas exchange from different individuals were collected and recut immediately under water. Furthermore, the cut ends of the branches were wrapped in damp towels, which were wrapped tightly by tiny plastic bags with some amount of water in it. Then, the branches were quickly placed in black plastic bags and brought back to the laboratory. All branches were recut underwater again, sealed with in black plastic bags, and allowed to rehydrate overnight in the laboratory. The cut ends of branches were wrapped by wax and parafilm (PM996; BEMIS). Leaf vulnerability curves were obtained using the rehydration method described by Brodribb & Holbrook (2003). According to the leaf vulnerability curve, we determined the maximum hydraulic leaf conductance (*K*_leaf_), the water potentials inducing 12% (*P*_12_) and 50% (*P*_50_) loss of the maximum conductance.

### Statistics and Graphics

Stomatal dynamics was fitted using a sigmoidal model in SigmaPlot 12.5, excluding stomatal transient WWR, and the time corresponding to decline of *g*_s_ by 10%, 50%, and 90% (*T*_10_‚ *T*_50_‚ *T*_90_) were obtained accordingly. The leaf vulnerability curves were fitted in SigmaPlot 12.5. Linear or nonlinear regression analyses were performed using species’ average values to examine the interspecific relationship between leaf water transport efficiency, gas exchange and water relations in nine mangrove species. Correlations were computed using mean values by performing linear or nonlinear regression analyses with stomatal dynamics parameters as dependent variables and stomatal morphology, leaf dehydration tolerance or leaf hydraulic capacitance as continuous independent variables across the nine mangrove species. The correlations were considered statistically significant if *P* < 0.05. All regression analyses and plots were performed using the R-project version 4.1.1 (https://www.r-project.org/).

## Results

### Stomatal dynamics is related to initial steady-state *g*_s_

The amplitude of variation in *g*_s_ response to a step-increase in VPD (Δ*g*_s_) was strongly related to *g*_s-initial_ (R^2^=0.993; Fig. 2a), but not to *g*_s-final_ (not shown). For other model parameters, λ was positively related to τ (R^2^=0.534; Fig. 2d) across the mangrove species, and the lower λ and τ were associated with greater *SL*_max_ (Fig. 2e-f) and faster closure speed in *g*_s_ (*T*_10_, *T*_50_, *T*_90_) in response to a step-rise in VPD (Fig. S2). Moreover, a significantly positive relationship between *g*_s-initial_ and λ existed for mangroves (R^2^=0.872; Fig. 2b). Correspondingly, a strong linear relationship between *g*_s-initial_ and time required to decrease *g*_s_ by 10%, 50% and 90% (*T*_10_, *T*_50_, *T*_90_) were found across the mangroves (Fig. S3). Meanwhile, the species with smaller Δ*g*_s_ had lower τ (Fig. 2c); however, neither λ nor τ showed significant relationships with *g*_s-final_ (not shown). These results indicate that initial steady-state *g*_s_ significantly influences the kinetics of stomatal response to a step-rise in VPD.

**Figure 2.**
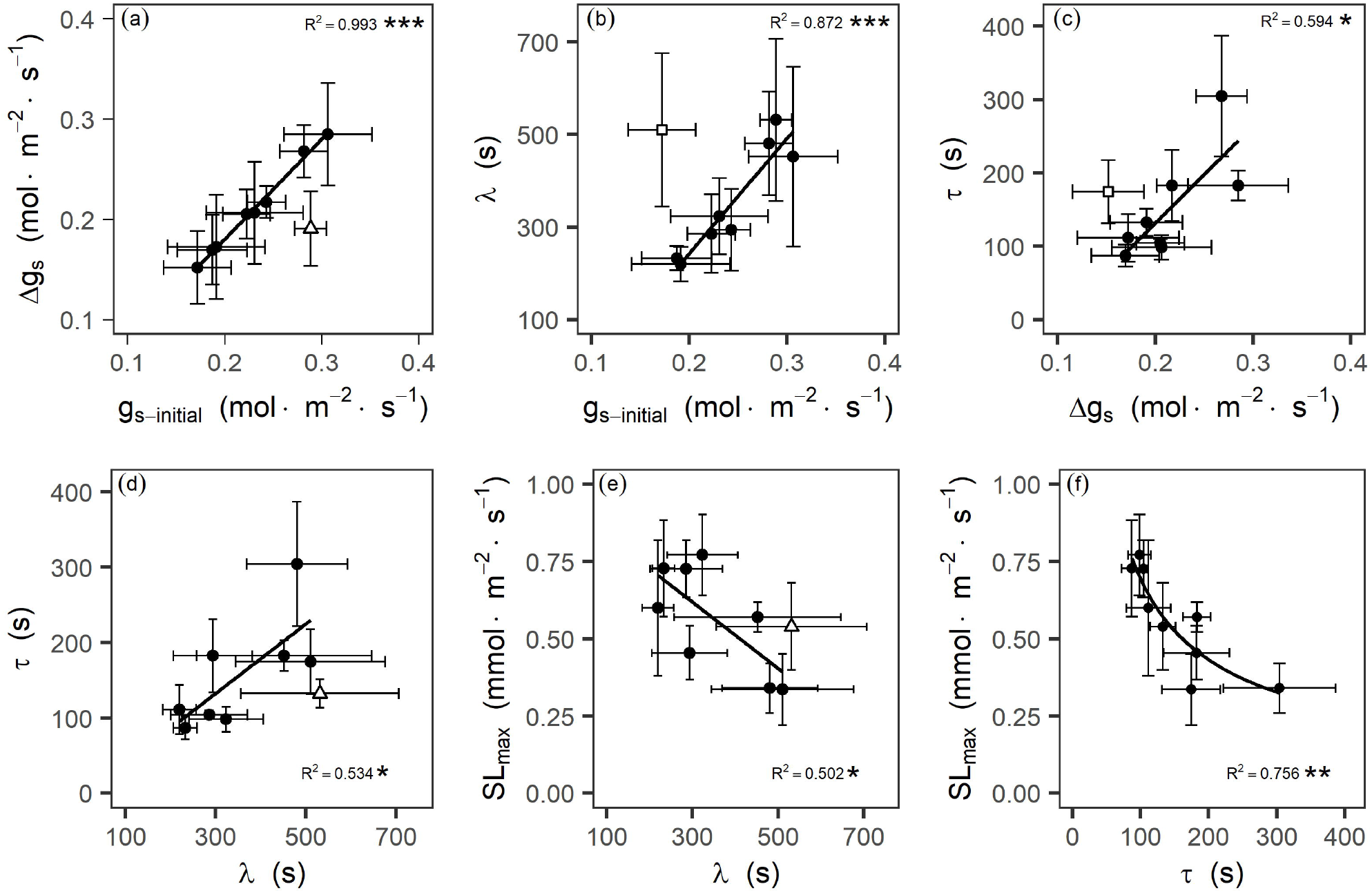
Relationships between parameters describing the kinetics of stomatal closure in response to temporal rising VPD. Correlations between the steady-state values of *g*_s_ at the initial step of the curve (*g*_s-initial_) and the amplitude of variation in *g*_s_ during an increase in VPD (Δ*g*_s_) (a), or the lag time λ for stomatal closing as a response to rising VPD (b). Correlations between the response time τ during stomatal closing response to a step change in VPD and Δ*g*_s_ (c), or λ (d). Non-linear relationship between the maximum speed of the stomatal VPD response at the inflection point of the curve (*SL*_max_) and λ (e), or τ (f). The regression lines were fitted for the significant relationships, excluding outliers indicated by white triangles (a, d-e) and white squares (b-c) are for *Avicennia marina* and *Rhizophora stylosa*, respectively; because there might be distinctive strategies in adaptation to drought for *Avicennia marina* with a trichome layer covering stomata on the abaxial leaf surface and *Rhizophora stylosa* with the most negative leaf *P*_50_ in nine mangrove trees. Points and error bars represent means and the standard error (±SE) for each species. * *p* < 0.05; \*\**p* < 0.01; \*\*\**p* < 0.001.

Furthermore, *g*_s-initial_, being close to actual maximum stomatal conductance (*g*_max_), was significantly and positively correlated with the maximum leaf hydraulic conductance (*K*_leaf_), the leaf osmotic potential at full turgor (π_o_), the theoretical maximum stomatal conductance (*G*_s-max_) among the mangrove species excluding an outlier, and *SD* was positively correlated with vein density (*VD*) and π_o_ (Fig. S4a-e). These data implied that *K*_leaf_, π_o_, *G*_s-max_ would influence the kinetics of stomatal response to a step-rise in VPD due to they had tight coordination with *g*_s-initial_. Moreover, the species with lower ε had higher *C*_leaf_ (Fig. S4f).

### Correlations between stomatal dynamics and morphology

The nine mangroves varied strongly in stomatal size (*SS*) and stomatal density (*SD*) (Fig. 3). These stomatal morphological traits were not related to stomatal dynamics in responses to a temporal rise in VPD across the nine mangroves (Fig. 3). However, for the five mangroves of *Rhizophoraceae*, τ was positively correlated with guard cell length (*GCL*) (R^2^=0.773; Fig. 3a) and negatively with *SD* (R^2^=0.78; Fig. 3b); and *SL*_max_ also showed a positive relationship with *SD* (R^2^=0.797; Fig. 3c). These results indicate that leaves with smaller and more numerous stomata have faster kinetics than larger stomata in response to a step-rise in VPD, but this is only valid across the closely-related species of the same family.

**Figure 3.**
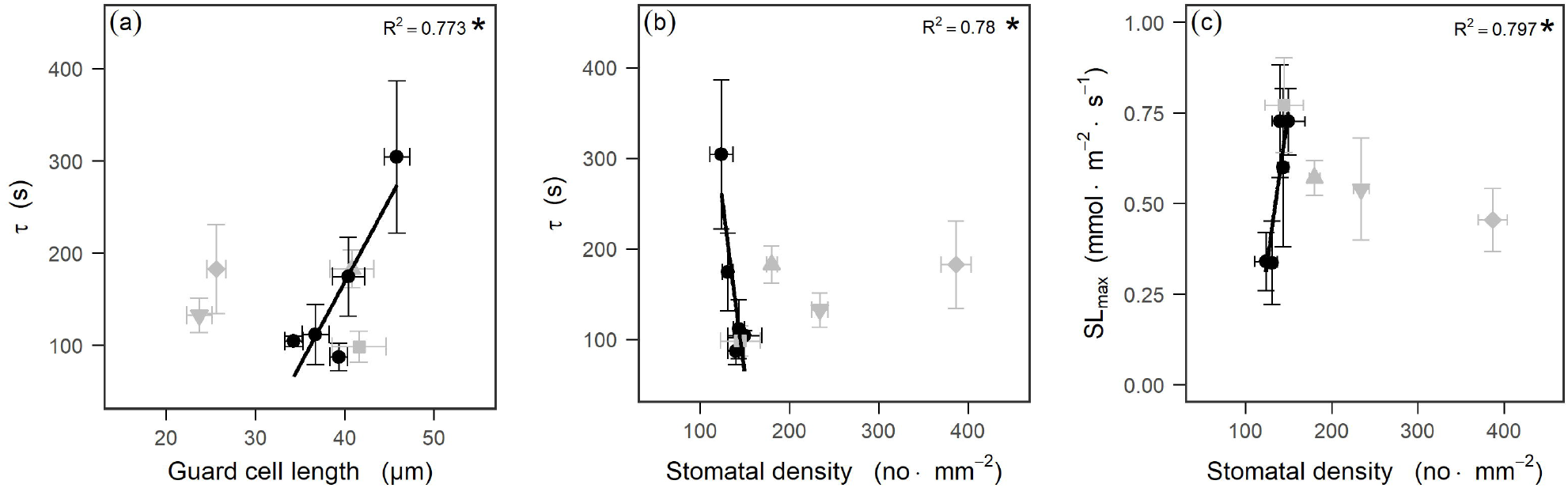
Relationships between parameters of the stomatal dynamics and stomatal morphology. Correlations between the response time τ during stomatal closing response to a step rising VPD and guard cell length (a), or stomatal density (b) across five mangrove species of *Rhizophoraceae*, between the maximum speed of the stomatal VPD response at the inflection point of the curve (*SL*_max_) and stomatal density (c) across the five species. The significant relationships were fitted with lines among five mangrove species of *Rhizophoraceae*, which is indicated by black circles. Grey symbols denote for the other four mangrove species of different family, which were not included in the regression line. There was no statistically significant relationship between parameters of the stomatal dynamics and stomatal morphology across the nine mangroves of various families (*p* > 0.05). Points and error bars represent means and the standard error (±SE) for each species, respectively. * *p* < 0.05.

### Stomatal dynamics in relation to leaf hydraulic traits

The nine mangrove species exhibited a two-fold variation in *P*_12_, ranging from - 1.07 MPa in *Aegiceras corniculatum* to -2.02 MPa in *Rhizophora stylosa* (Fig. 4). The species with less negative *P*_12_ had higher *SL*_max_ and faster τ and λ (Fig. 4), but *P*_50_ was not statistically related to *SL*_max_, τ, and λ (not shown), indicating that *P*_12_ and not *P*_50_, might be involved in stomatal regulation and that a lower vulnerability of leaves to dehydration could lengthen stomatal closing time in response to high VPD. *C*_leaf_ was positively and significantly correlated (R^2^=0.775) with τ and was negatively correlated (R^2^=0.714) with *SL*_max_ (Fig. 5), indicating that under a high transpiration demand, high *C*_leaf_ could delay the stomatal closure due to buffering a sudden drop in water potential of leaf tissues.

**Figure 4.**
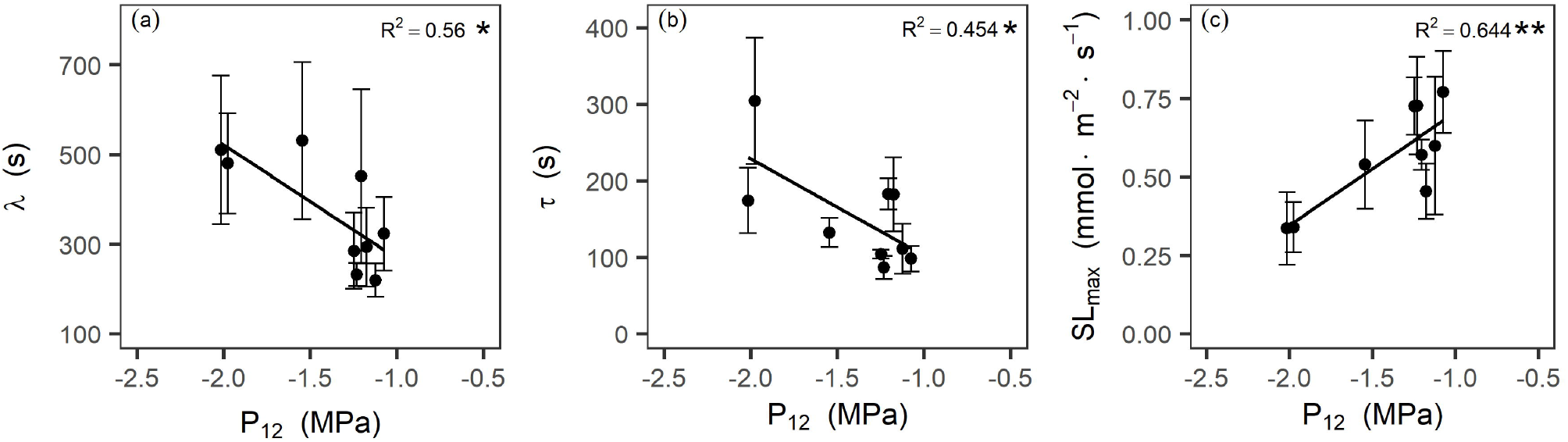
Relationships between hydraulic vulnerability to dehydration of leaves and parameters describing the kinetics of stomatal closure in response to rising VPD. Correlations between the water potential inducing 12% (*P*_12_) loss of the maximum conductance of leaves and the lag time λ for stomatal closing as a response to VPD (a), the response time τ during stomatal closing response to a step increase in VPD (b) and the maximum speed (*SL*_max_) of the stomatal VPD response at the inflection point of the curve (c). Points and error bars represent means and the standard error (±SE) for each species, respectively. \**p* < 0.05; \*\**p* < 0.01.

**Figure 5.**
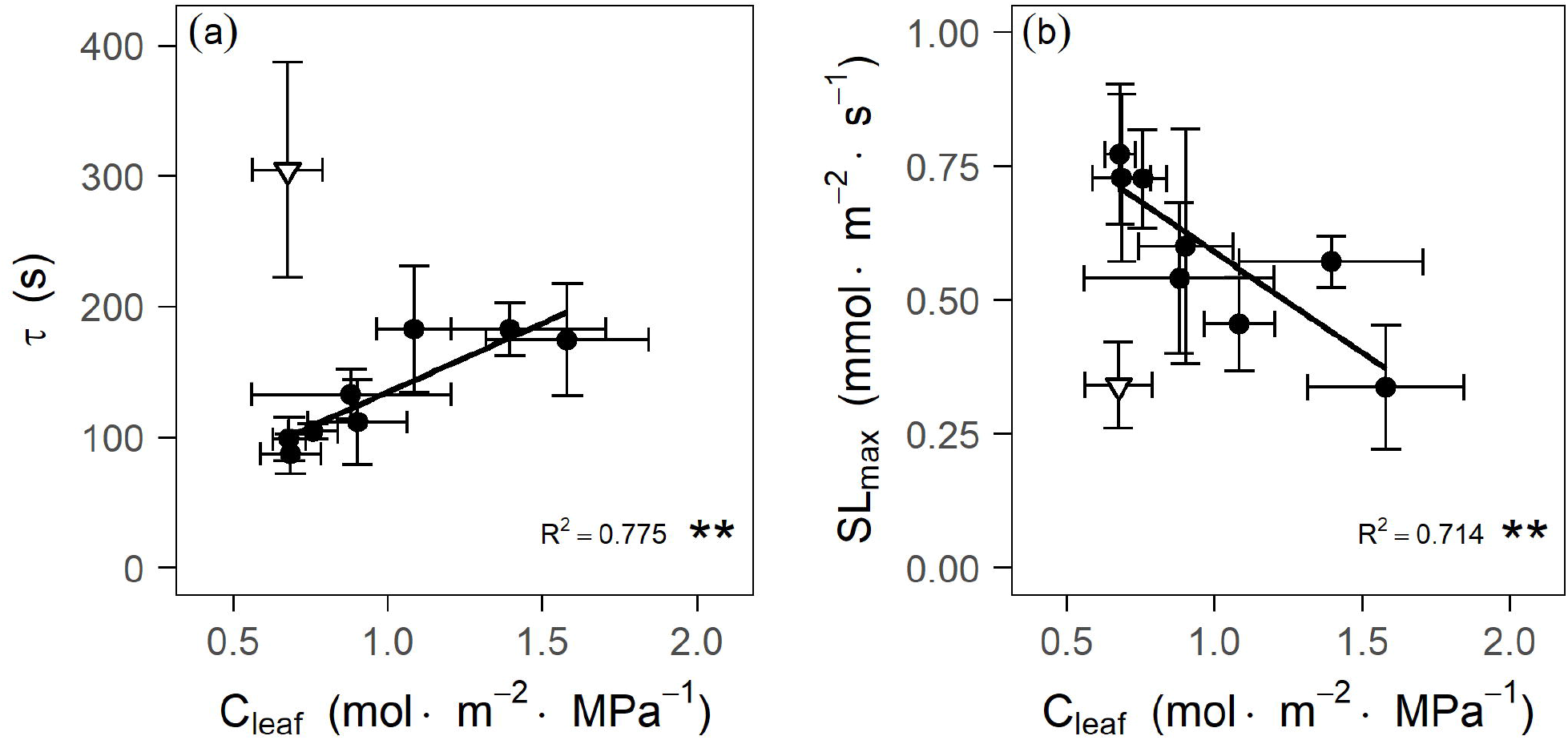
Relationships between leaf hydraulic capacitance (*C*_leaf_) and parameters describing the kinetics of stomatal closure in response to rising VPD: the response time τ during stomatal closing response to a step-rise in VPD (a), and the maximum speed (*SL*_max_) of the stomatal response at the inflection point of the curve (b). White inverted triangle (a-b) is for *Ceriops tagal* with the most negative π_tlp_, which showed an extremely strong the ability of maintaining leaf osmotic pressure and could be distinguished from the nine mangrove species; it thus was excluded in the correlations. Points and error bars represent means and the standard error (±SE) for each species, respectively. \*\**p* < 0.01.

## Discussion

We found strong evidence that the morphology of stomata and hydraulic traits of leaves influences stomatal response to a sudden increase in VPD in mangrove trees. In our transient VPD perturbation experiment, we observed that *P*_12_, not *P*_50_, significantly controlled stomatal dynamics and also higher *C*_leaf_ can delay stomatal closure. *P*_12_ as a strong regulator of kinetics of stomatal closure could provide a timely protection of the leaf hydraulic system when facing atmospheric drought. This regulatory behavior of stomata could be of vital importance for the long-term survival of mangrove trees in a stressful environment.

### Effects of maximum stomatal conductance and stomatal morphology on stomatal dynamics

In this study, we assessed the kinetics of stomatal response to a step-rise to a high VPD among nine mangrove species *in situ*. We found *g*_s-initial_, not *g*_s-final_, determined Δ*g*_s_. Moreover, we observed a tendency of increasing λ and τ with higher *g*_s-initial_ and that the lower λ and τ were associated with greater *SL*_max_. A higher *g*_s-initial_, λ and τ could help plants achieve greater carbon gain (Prentice *et al*. 2014; Wolf *et al*. 2016; Anderegg *et al*., 2018). However, this would result in a greater water potential gradient from the petiole and vein xylem to the evaporating site below the stomata immediately after a step-rise in VPD (Sperry, 2000; Buckley *et al*., 2011), and thus higher hydraulic risk. This lower water potential could drive greater ABA biosynthesis in these plants after a step change in VPD and lead to the higher λ and τ. Plants with a more sensitive stomatal response with a steep *SL*_max_ to rapidly decrease *g*_s_ to the new steady-state *g*_s-final_ for avoiding desiccation (Fig. 2e-f). Overall, *g*_s-initial_ strongly influenced stomatal behavior after a step-rise in VPD. In fact, *g*_s-initial_ is close to actual maximum stomatal conductance (*g*_max_), thus we hypothesized that traits that determine *g*_*max*_ should also influence the kinetics of stomatal response to a step-rise in VPD. Firstly, we found that the osmotic pressure at the full turgor (π_0_) was an emergent determinant of gas exchange and its regulation behavior. This finding is consistent with the recent proposal that species with higher π_o_ and π_tlp_ had higher *g*_max_ and greater sensitivity in stomatal closure during leaf dehydration over a wide range of species (Henry *et al*., 2019). Our work on the nine mangroves, which have more negative π_o_ and π_tlp_ than most of terrestrial plants, provides support for the idea of stomatal safety-efficiency trade-off (Henry *et al*., 2019). Secondly, our work shows that mangroves have a functional coordination between water transport capacity and maximum rates of gas exchange, enabling the water balance between hydraulic supply and demand.

We also observed that, a significant linear relationship between theoretical maximum stomatal conductance (*G*_s-max_) and *g*_s-initial_, and stomatal density (*SD*) was related with π_o_ and vein density (*VD*) across the nine mangrove species. These reflect a potential mechanistic control of stomatal size and density on stomatal responses to environmental fluctuations. Smaller, denser stomata have often considered to have faster kinetics (Drake *et al*., 2013), which is because changes in solutes can be mechanically facilitated owing to a greater surface area to volume ratio of the guard cell membrane in smaller than larger stomata (Drake *et al*., 2013; Raven, 2014; Durand *et al*., 2019). In our study, we observed a trend of increasing speed of closure with increasing *SD* and decreasing guard cell length (*GCL*) in the five mangrove species of *Rhizophoraceae*, but this pattern was not found in nine mangrove species of various families. Elliott-Kingston *et al*. (2016) found no relationship between closure rapidity and size or density of stomata from studies of stomatal kinetics along an evolutionary diverse series of species (including fern, cycad, conifers and angiosperms). Likewise, McAusland *et al*. (2016) examined rapidity of stomatal closure from species over a range of crops with differences in stomatal morphology (kidney- or elliptical-shaped) and found that stomatal kinetics was also not explained by the size of stomata. Overall, our study is generally consistent with previous studies and suggests that the relationship between stomatal closing speed in response to a step-rise in VPD and stomata size is conserved in closely related species. There may be mechanisms that affect stomatal movement speed in response to ambient environmental fluctuation independent of size across species with diverse stomata morphology. Gérardin *et al*. (2018) found that in *Nicotiana tabacum* grown under shade or drought conditions, the number of epidermal cells around guard cells could influence stomatal response speed; and stomata with a few peripheral epidermal cells could undergo a lower extrusion force from epidermal cell turgor than those with a higher density of peripheral epidermal cells. Furthermore, guard cells with subsidiary cells might allow stomatal complexes to overcome an epidermal mechanical effect by a reciprocal exchange of potassium between the two cells (Gray *et al*., 2020). The cytoskeleton (Higaki *et al*., 2010; Eisinger *et al*., 2012) and cell wall elasticity (Carter *et al*., 2017; Woolfenden *et al*., 2017) have also been proposed to be partially involved in changes in guard cell shape and volume, which are the basic sources of stomatal movement. Recently, Jalakas *et al*. (2021) discussed the molecular mechanisms of stomatal closure in response to a sudden increase in VPD and suggested that different ABA signaling pathways could have different contributions to VPD-induced stomatal closure of angiosperms. Lastly, hydraulic signals are the basic feedback mechanisms on controlling stomatal movements (Buckley & Mott, 2002; Buckley, 2005; Franks, 2013; Brodribb *et al*., 2014; Tardieu, 2016; Buckley, 2019), as shown by the present study that stomatal behavior was strongly regulated by hydraulic messages (e.g., the water potential inducing 12% loss of leaf hydraulic conductance, *P*_12_) and water-relations traits of leaves (discussed below).

### Leaf hydraulic vulnerability and capacity affect stomatal regulation behavior

Timely stomatal closure helps plants avoid death during a drought (Martin-StPaul *et al*., 2017; Hochberg *et al*., 2017; Li *et al*., 2018), at the cost of a cessation of photosynthesis and potential transpirational cooling of leaves (Flexas & Medrano, 2002; Leigh *et al*., 2017), having a negative effect on plant growth (Mitchell *et al*., 2013; Sevanto *et al*., 2014). Our results indicated that *P*_12_ is a strong regulator of the kinetics of stomatal closure in response to a temporal rise in VPD in the mangroves. Firstly, *P*_12_, which is the initial inflection point of the vulnerability of leaf hydraulic conductance to drought, is considered as a potential trigger for stomatal closure as explained by simple hydraulic feedback (Raschke & Khül, 1969; Buckley & Mott, 2002; Buckley, 2005). Secondly, *P*_12_ reflects the primary expansion of hydraulic resistance to bulk flow of the liquid-phase moving from xylem into the bundle sheath to epidermal cells and guard cells within leaves during dehydration (Sack & Holbrook, 2006). It drives an initial increase in the back pressure of the epidermal pavement cells on guard cells (Darwin, 1898; Cowan, 1977), triggering hydraulic feedback to overcome the epidermal mechanical advantage for stomatal closure (Buckley, 2019). Lastly, the whole-leaf hydraulic resistance also includes the vapour-phase diffusion from the evaporating sites to stomata (Buckley *et al*., 2017), which is linked with the liquid transport in transpiring leaves and closely related to water potential (Rockwell *et al*., 2014). It, therefore, seems likely that species with more negative *P*_12_ have greater vapour diffusion through stomata (Peak & Mott, 2011), which would delay stomatal closure in response to atmospheric drought. However, the water potential corresponding to 50% loss in leaf hydraulic conductance (*P*_50_) was not significantly correlated with the kinetics of stomatal closure in response to a step-rise in VPD (not shown) across mangroves. *P*_50_ reflects the long-term adaptive abilities of plants to deal with a drought event, while *P*_12_ reflects the short-term regulatory strategy of plants to instantaneously optimize the marginal benefit in response to any hydraulic perturbation (Sperry, 2000; Brodribb *et al*., 2016; Choat *et al*., 2018; Zhang *et al*., 2020). The adaptation of *P*_12_ as a strong regulator of kinetics of stomatal closure could timely prevent a dramatic decline in leaf water potential and avoid xylem embolism, and thus is of crucial importance for the long-term survival of mangrove trees.

Additionally, we observed that mangroves developed thick or even upper and lower hypodermis, contributing a large proportion of lamina thickness and storing a considerable amount of water inside leaves (Nguyen *et al*., 2017a; Nguyen *et al*., 2017b), thus had high *C*_leaf_. Leaves with a high *C*_leaf_ can buffer fluctuation in leaf water potentials when facing atmospheric drought, by compensating transpirational water loss from storage water. Therefore, we observed a significant trend of increasing time of stomatal closure in response to a step-rise in VPD in the mangrove species with higher *C*_leaf_. This is consistent with previous studies (Scholz *et al*., 2011; Martins *et al*., 2016; Fu *et al*., 2019; Xiong & Nadal, 2020). Although high *C*_leaf_ of the mangroves may delay stomatal closure at high VPD, species with a high *C*_leaf_ had a lower overall *T*_50_ values than the six woody species mentioned by Fu *et al*. (2019). Thin leaves of the six woody species examined by Fu *et al*. (2019), could systematically alleviate the instantaneous decline of leaf water potential under an atmospheric drought due to a shorter hydraulic pathway from vascular bundle to stomata so that generate a smaller water potential gradient across mesophyll tissues (Brodribb *et al*., 2007; de Boer *et al*., 2016; Buckley *et al*., 2017) and thus higher *T*_50_, because water potential would not drop close to π_tlp_ at which point ABA would be synthesized in leaves (McAdam & Brodribb, 2016). By contrast, the thick leaves of mangrove trees with longer outside-xylem pathways of water transport are difficult to maintain water balance between mesophyll cells when facing rapid changes in atmospheric moisture (Brodribb *et al*., 2007; de Boer *et al*., 2016; Scoffoni *et al*., 2017), resulting in a larger change in turgor pressure of guard cells. This might accelerate the transmission of hormonal signal about ABA biosynthesis inside leaves for faster stomatal closure to reduce water loss (Franks, 2013; Tardieu, 2016; McAdam & Brodribb, 2016; Buckley, 2019) and thus, for mangrove trees, lower *T*_50_. Moreover, epidermal cells of mangroves are smaller than other angiosperms and also smaller than their own stomata (Jiang *et al*., 2023) meaning that stomata of mangroves have a higher density of peripheral epidermal cells. This design feature might contribute to the stomatal movement as suggested by Gérardin *et al*. (2018), and thus faster stomatal closure for mangroves. Furthermore, mangroves for the adaptation of high salinity of intertidal habitats have evolved a series of water-protection traits, such as highly embolism-resistant xylem (Jiang *et al*., 2017; Jiang *et al*., 2021; Jiang *et al*., 2022), high hydraulic-capacity in leaf and root tissues (Aritsara *et al*., 2022), and extreme osmotic adjustment (Reef & Lovelock, 2015). They therefore may have enhanced stomatal regulation with a highly sensitive stomatal response to VPD suited to this adaptation to water-deficit prone habitats.

Our results also showed that the leaves of mangroves had a high bulk modulus of elasticity (ε). This means that cell walls are rigid, and could provide mechanical strength and maintain cellular hydration during dehydration. Our results also showed a negative relationship between *C*_leaf_ and ε across the mangrove species. This finding was consistent with recent framework on water movements and storage dynamics (Xiong & Nadal, 2020), suggesting that a tight coordination between capacitance and resistance of deformation and shrinkage under tension.

Notably, the three distinctive strategies in adaptation to drought can be distinguished from the nine mangrove species, as represented by *Ceriops tagal, Rhizophora stylosa* and *Avicennia marina*, respectively. *Ceriops tagal* had the lowest π_tlp_ (most negative value), showed an extremely strong the ability of maintaining leaf osmotic pressure. *Rhizophora stylosa* had the most negative leaf *P*_50_, i.e., the most resistance to dehydration in leaf hydraulic conductance. *Avicennia marina* has a trichome layer covering stomata on the abaxial leaf surface, which can effectively reduce transpirational water loss by increasing the boundary-layer resistance to vapor diffusion. The functional traits of the nine mangroves have acted excellently in coping with environmental stress, but they may loosen the linking stomatal behaviors with leaf hydraulic capacities and water relations at some point in the regulation chain.

## Conclusions

We here show that stomatal movement in response to a temporal rise in VPD, stomatal morphology and hydraulic traits, in nine true mangrove species, are coordinated. Smaller, denser stomata had faster, stronger kinetics of stomatal response in closely related species. Water-protection traits, such as *C*_leaf_ and *P*_12_ were negatively associated with stomatal closure speed when facing atmospheric water deficit. Our work provides a quantitative framework to better understand stomatal regulation behavior of mangroves in a stressful environment. Further studies need to quantify the contribution of variation and dynamics in both leaf hydraulic conductance and leaf capacitance to the stepwise regulation of stomatal dynamics in response to any environmental perturbation that changes water status in the plant.

## Acknowledgements

We acknowledge the collaboration of the Dongzhaigang National Nature Reserve and Qinglan Port Provincial Mangrove Nature Reserve, and the Forestry Department of the Hainan Province for permission to conduct the experiment in the mangrove reserves. The authors thank Prof. Junjie Zhu for the experimental design, Yang Wei for field assistance, Jin-Yan Lei and Pei-Xin Cui for their help in leaf anatomy.

## Supporting Information

**Fig. S1** Stomatal responses to a step-rise in VPD of the mangrove plants. Responses normalised (a) to the average values of observed *g*_s_ and responses relativized (b) to the steady-state values of calculated *g*_s_ at the start of curve were presented on the time course after a step increase in VPD from ∼0.95 to ∼2.65 kPa at t = 0 min. Brown, *Sonneratia alba*; orange, *Avicennia marina*; turquoise, *Ceriops tagal*; skyblue, *Xylocarpus granatum*; pink, *Aegiceras corniculatum*; lightblue, *Bruguiera gymnorhiza*; green, *Bruguiera sexangula*; purple, *Kandelia obovate*, olive, *Rhizophora stylosa*.

**Fig. S2** Relationships between parameters of the stomatal dynamics and the timing of changes in *g*_s_ in response to a single step-rising vapour pressure deficit (VPD). Correlations between the time to decline of *g*_s_ by 10% (*T*_10_) and the lag time λ for stomatal closing as a response to step-rise in VPD (a), the response time τ during the stomatal closing response (b), and the maximum speed (*SL*_max_) of the stomatal response at the inflection point of the curve (c). Correlations between the time to decline of *g*_s_ by 50% (*T*_50_) and λ (d), τ (e), and *SL*_max_ (f). Correlations between the time to decline of *g*_s_ by 90% (*T*_90_) and λ (g), τ (h), and *SL*_max_ (k). Points and error bars represent means and the standard error (±SE) for each species, respectively. \**p* < 0.05; \*\**p* < 0.01; \*\*\**p* < 0.001.

**Fig. S3** Relationships between gas exchange and the timing of changes in *g*_s_ in response to a single step-rising vapour pressure deficit (VPD). Correlations between initial *g*_s_ (*g*_s-initial_) and the time corresponding to decline of *g*_s_ by 10%, 50%, and 90% (a, *T*_10_; b, *T*_50_; c, *T*_90_), respectively. White squares (a-c) denote the outlier *Rhizophora stylosa*, which were excluded in the correlations. Points and error bars represent means and the standard error (±SE) for each species, respectively. \**p* < 0.05; \*\**p* < 0.01; \*\*\**p* < 0.001.

**Fig. S4** Relationships between gas exchange and water-related traits. Correlations between initial *g*_s_ (*g*_s-initial_) and the maximum leaf hydraulic conductance (*K*_leaf_) (a), the leaf osmotic potential at full turgor (π_o_) (b), the theoretical maximum stomatal conductance (*G*_s-max_) (c) across mangroves, respectively. Correlations between Log_10_-transformed values of stomatal density (*SD*) and vein density (*VD*) (d), log10-transformed values of π_o_ (e) across mangroves, respectively. Correlations between leaf capacitance (*C*_leaf_) and bulk modulus of elasticity (ε) (f) across mangroves. Note the regression line fitted without the outlier denoted by a white point in **b** and **c**, which is *Ceriops tagal* and *Xylocarpus granatum*, respectively. Points and error bars represent means and the standard error (±SE) for each species, respectively. \**p* < 0.05; \*\**p* < 0.01.

