## Supplementary figures and images for "Stomatal dynamics are regulated by leaf hydraulic traits and guard cell anatomy in nine true mangrove species"

### Supplemental Fig. S1

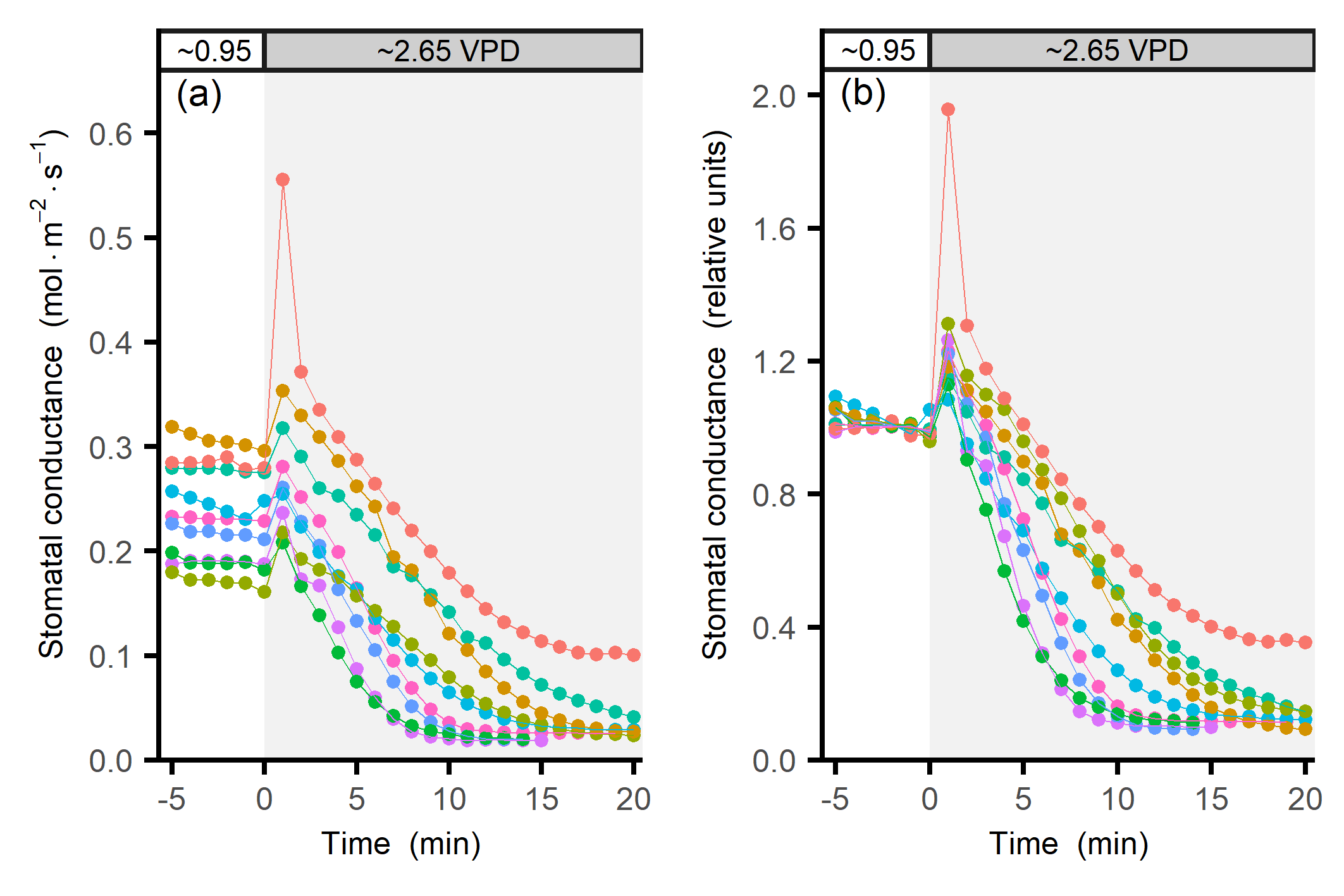

### Supplemental Fig. S2

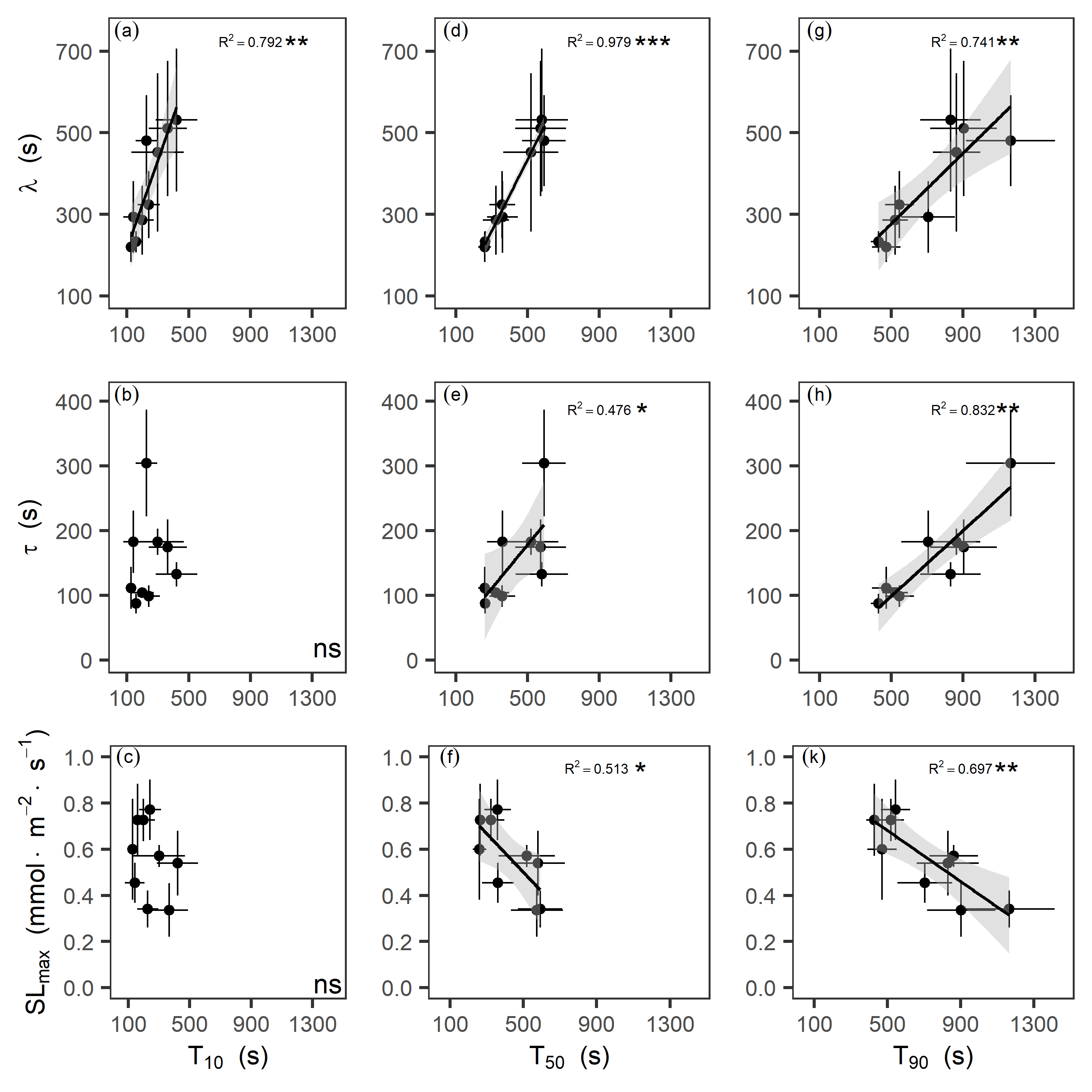

### Supplemental Fig. S3

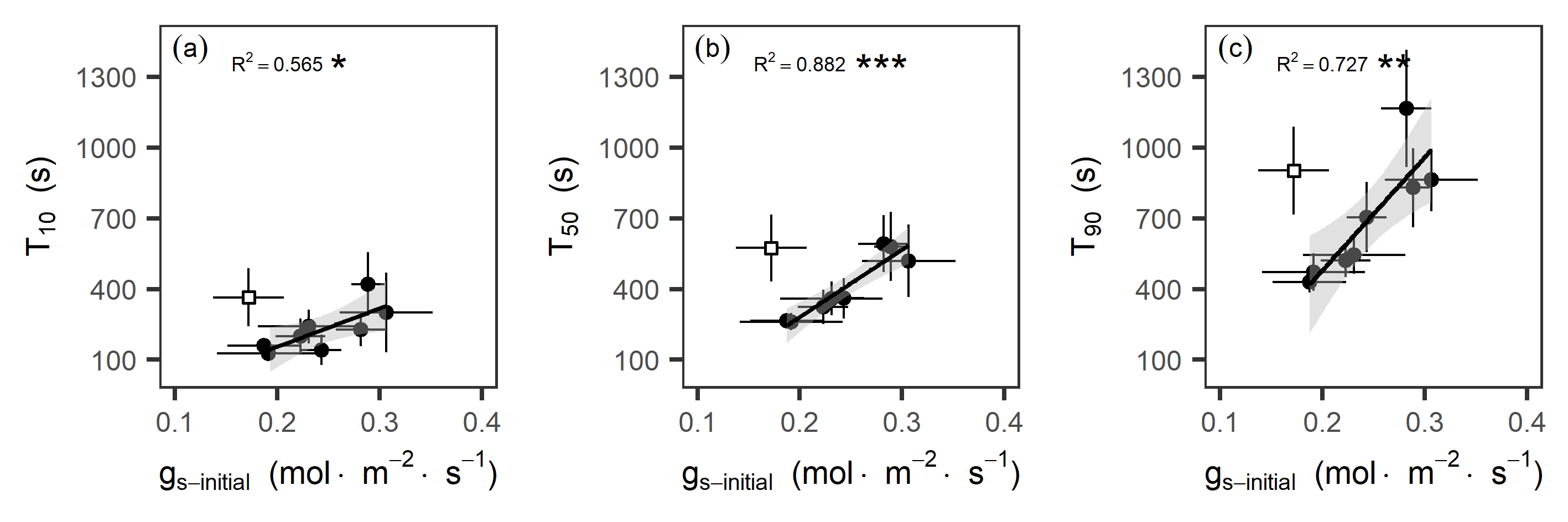

### Supplemental Fig. S4

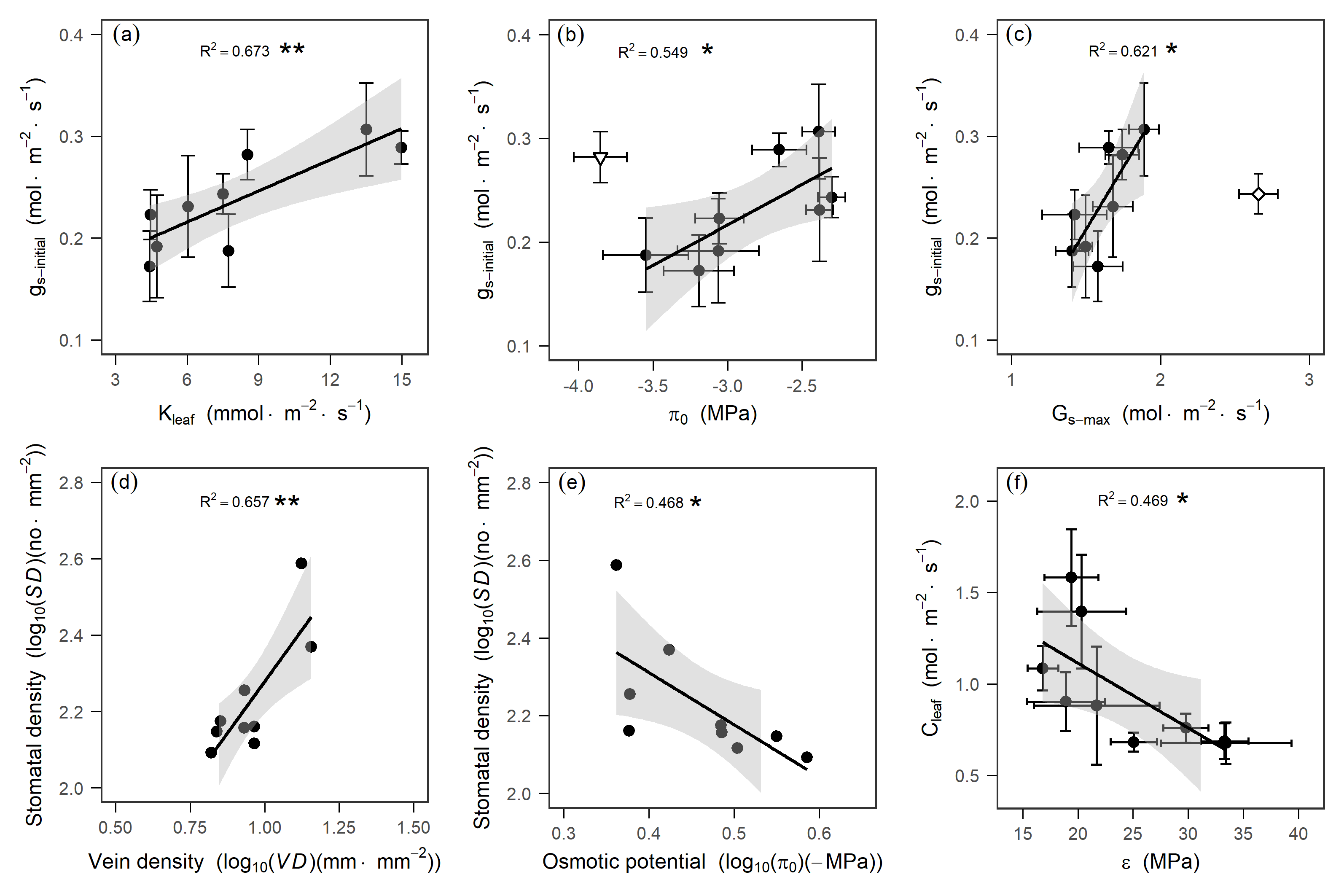
